# Concerted remodelling of the postsynaptic spine and RNA granule by cLTP

**DOI:** 10.1101/2025.07.16.665171

**Authors:** Mònica B. Mendoza, Galal Yahya, Kimberly Alomoto, Adriana Soria-Villalba, Martí Aldea, Carme Gallego

## Abstract

Synaptic plasticity is the cellular foundation of learning and memory and, for these plastic changes to be stabilised into long-term memory, proteins must be synthesised at the synapse^1,2^. RNA granules ensure that specific mRNAs are delivered and translated at the right time and place^3,4^, but the signalling mechanisms linking synaptic activation to local translation are still largely unknown. Here, to investigate how postsynaptic signals modulate the RNA granule, we employed a spatially-restricted biotinylation approach to quantify protein accessibility and proximity in the postsynaptic and RNA granule subproteomes. Upon chemical long-term potentiation (cLTP), we observed a sharp increase in the accessibility of ribosomes, translation factors and RNA binding proteins belonging to the RNA granule. Similarly, strong alterations were observed in proteins of the postsynaptic density, but mostly in those involved in signalling. Specific proximity to DBN1 and IGF2BP1 as postsynaptic and RNA granule reporters unveiled a shift of the translation machinery towards the postsynaptic compartment, whereas specific translation initiation factors and RNA helicases displayed marked changes in their proximity to DBN1 and/or IGF2BP1. Finally, alteration of synapse development and signalling by DBN1 downregulation caused a highly significant decrease in the proximity of RNA granule proteins to IGF2BP1 after synaptic stimulation, establishing a causal link between postsynaptic events and RNA granule dynamics.

## Introduction

Synaptic plasticity, which forms the basis of cognitive function and memory, is the ability of synapses to modify their strength in response to stimulation, orchestrated by tightly regulated molecular remodelling within dendritic spines^5,6^. Increasing evidence in recent years indicates that synapses are highly compartmentalised^7^. The organisation of molecular complexes into condensed molecular networks by phase separation allows the autonomous formation of distinct functional compartments on both sides of synapses. On the postsynaptic side, underneath the postsynaptic plasma membrane is a highly electron-dense zone known as the postsynaptic density (PSD), composed of a densely packed network of receptors, scaffold proteins, and signalling molecules. The PSD appears to be further compartmentalised with a more electron-dense layer known as the core lying immediately beneath the postsynaptic membrane and the pallium, a relatively less electron-dense layer facing the spine soluble phase^8,9^. Such highly compartmentalised organisation of the synapse is not static. Instead, molecular constituents can exchange between postsynaptic subcompartments in response to synaptic stimulation. For example, postsynaptic spines are known to enlarge during long-term potentiation (LTP), at the same time that a reorganisation of the actin cytoskeleton occurs^10,11^. In cultured hippocampal neurons, the postsynaptic pallium becomes more electron-dense after depolarisation with KCl or treatment with glutamate, likely due to the accumulation and redistribution of postsynaptic proteins^12^. Furthermore, long-term stabilisation of the extended spine requires synthesis of synaptic proteins, stabilisation of actin filaments and enlargement of the postsynaptic compartment^13^.

In addition to postsynaptic condensates in dendritic spines, dendritic shafts contain neuronal RNA granules, biomolecular condensates composed of RNA-binding proteins (RBPs) and mRNA molecules. These granules are dynamic assemblies and have been implicated in mRNA transport and translation regulation^1,3,14^. A number of studies indicate that the granules consist of stably paused (stalled) ribosomes^15–18^ where translation of the stored mRNA resumes upon synaptic stimulation^2,19^.

Although numerous components of the postsynaptic compartment and RNA granules are already known from proteomics and transcriptomics studies^15,20–25^, how such different membraneless compartments communicate and coordinate with each other in regulating synaptic development and signalling is poorly understood.

## Results

### cLTP alters protein accessibility

Proximity labelling has been applied to study a paradigmatic biomolecular condensate by fusing stress-granule protein components to APEX2, an engineered enzyme that catalyses the addition of biotin to proximal proteins within a very small diffusion radius in a native context^26,27^. However, protein-protein interactions found in these biomolecular condensates may also be detected in the non-condensed state, making a precise quantitative analysis of the condensation process *in vivo* difficult. As a completely different approach, we reasoned that a free cytoplasmic form of APEX2 would add biotin to proteins as a direct function of their accessibility, thus reporting on their condensation status *in vivo* with high temporal resolution (Fig. 1a). Moreover, since as much as 99% of the cytoplasmic proteome of a single neuron resides in neurites^1^, we assumed that our approach would particularly report on the composition and dynamics of proteins in the dendritic arbour, including postsynaptic spines and RNA granules. Thus, we expressed APEX2 (which contains a constitutive nuclear export signal) from a lentiviral vector in mouse cortical neurons and performed biotinylation assays at 17DIV. Prior to the addition of hydrogen peroxide, neurons were exposed to biotin for 30 min (basal), subject to chemical long-term potentiation (cLTP) for 10 min and returned to normal medium for 30 min (post-cLTP) (Extended Data Fig. 1a). Thanks to the nuclear export signal, the APEX2 protein displayed a mostly cytoplasmic distribution in cortical neurons (Extended Data Fig. 1b). Overall, we quantified 2560 proteins in biotinylated pulldowns from cortical neurons expressing APEX2 and, as expected from a large diversity in protein accessibility, levels in streptavidin beads and input samples correlated only to a minor extent (Extended Data Fig. 1c). Moreover, biotinylation efficiency was not obviously affected by the predicted number of tyrosines on the surface of each protein (Extended Data Fig. 1d). No cell components were significantly enriched in highly biotinylated proteins but, as expected from their inherently low accessibility in the cytoplasm, proteins belonging to the nucleus, mitochondria and the lumen of vesicles or the ER displayed very low biotinylation efficiencies (Extended Data Fig. 1e). Notably, protein components of the RNA granule and the ribosome also exhibited a very low biotinylation efficiency under basal conditions. In all, our data support the notion that the efficiency of biotinylation by a free cytoplasmic APEX2 enzyme, as a reporter of accessibility, can be used to quantify changes in protein assemblies such as biomolecular condensates.

**Fig. 1.**
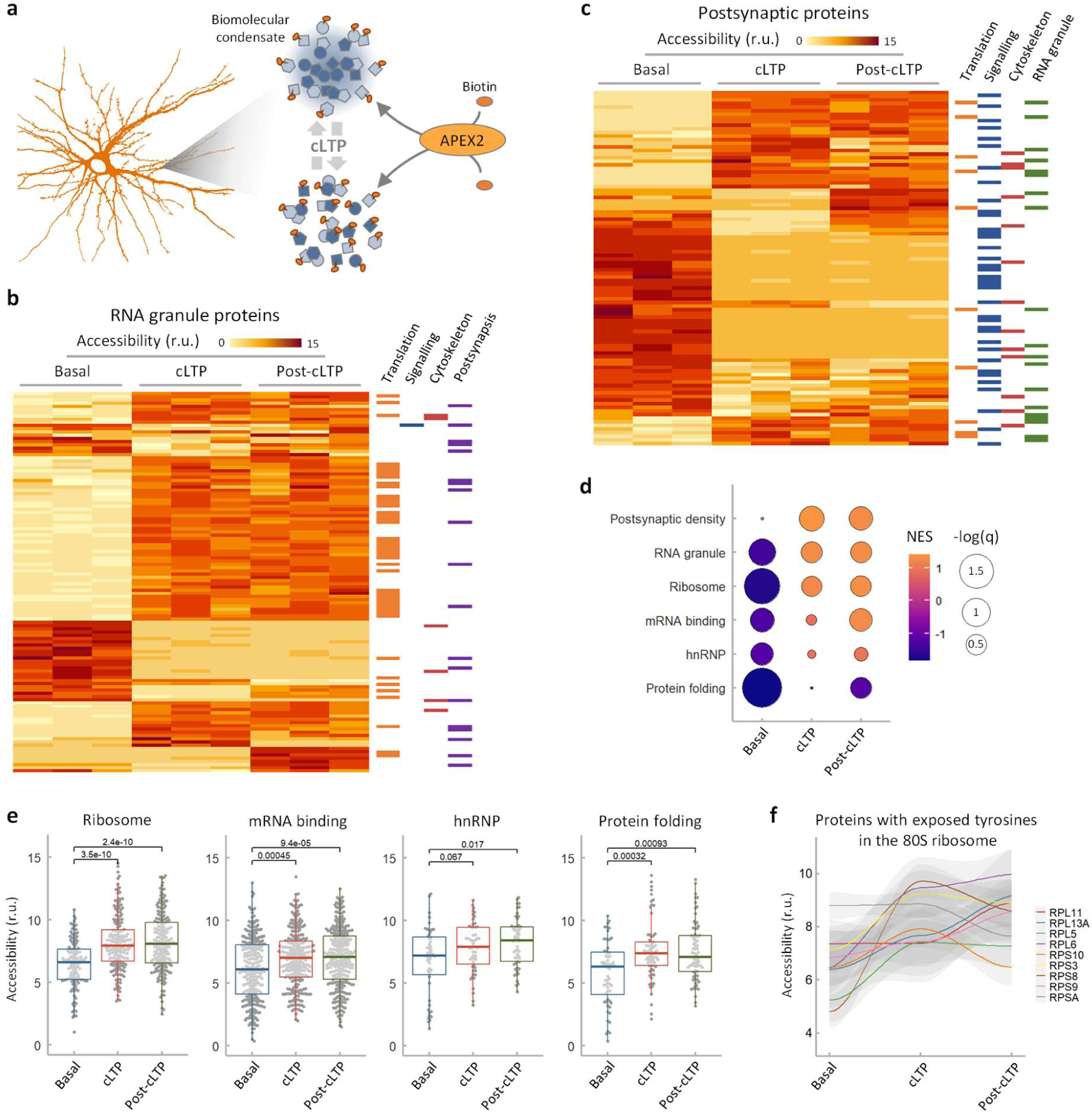
cLTP alters accessibility of postsynaptic and RNA granule proteins. **a**, Schematic of the experimental approach to analyse changes in protein accessibility in cortical neurons during cLTP. **b**,**c**, Accessibility scores under basal, cLTP and post-cLTP conditions of RNA granule (b) and postsynaptic (c) proteins. Shown are three replicate values for proteins displaying at least a 10-fold change in accessibility in any condition. Relevant protein categories are also indicated. **d**, GSEA results for the indicated selection of protein sets. **e**,**f**, Accessibility scores corresponding to the indicated protein sets (e) or individual proteins (f) under basal, cLTP and post-cLTP conditions.

Cortical neurons subject to cLTP showed a remarkable change in the accessibility of proteins belonging to the RNA granule and postsynaptic compartment (Fig. 1b,c). Of note, changes in protein accessibility were usually sustained during the post-cLTP period. The vast majority of proteins with altered accessibility in the postsynaptic compartment are involved in signalling tasks. Regarding the RNA granule, most proteins with increased accessibility participate in translation. Among them, ribosomal proteins, mRNA-binding proteins and, specifically, hnRNPs were strongly represented (Fig. 1d), and displayed highly significant increases in accessibility under cLTP and post-cLTP conditions (Fig. 1e).

Finally, cLTP produced an increase in accessibility of protein chaperones that partially reverted later on (Fig. 1d,e). Although biotinylation efficiency under basal conditions partly correlated with the number of exposed tyrosines on the surface of ribosomal proteins (Extended Data Fig. 1f), the increase in their accessibility upon cLTP did not obviously depend on the number of hidden tyrosines in the 80S ribosome (Extended Data Fig. 1g), suggesting that the observed increase in accessibility is not due to ribosome dissociation into single components. The total level of ribosomal proteins was barely affected by cLTP (Extended Data Fig. 1h), and almost all ribosomal proteins with tyrosines exposed on the surface of the 80S ribosome became more biotinylated upon cLTP (Fig. 1f), which indicated an increase in accessibility of the ribosome itself.

### Dendritic spine and RNA granule location

Synaptic plasticity has been shown to involve remodelling of the actin cytoskeleton ^28,29^ and increased local translation of the actin mRNA, which is recruited to the RNA granule by the mRNA-binding protein IGF2BP1^3^. On the other hand, DBN1 is among the actin-binding signalling proteins with the highest intrinsic disorder score (Extended Data Fig. 2a,b). In addition, DBN1 is subject to extensive phosphorylation (Extended Data Fig. 2c) and specifically accumulates in dendritic spines to coordinate actin and microtubules in neuronal cells^30^. Thus, we chose DBN1 and IGF2BP1 to test whether localization of IGF2BP1-containing RNA granules and synaptic spines show a correlation in dendrites, and expressed the corresponding fusions to compatible fluorescent proteins with a neuritic tracker in hippocampal neurons (Fig. 2a). More specifically, we measured the distance between the postsynaptic spine and the nearest RNA granule (Fig. 2b) and plotted these data as a function of spine size or DBN1 level in spine. Interestingly, spine-RNA granule distance displayed a sigmoidal dependence on spine size, indicating that at least a fraction of RNA granules localises to the dendritic shaft of spines above a critical size (Fig. 2c). Levels of HOMER1, a postsynaptic scaffolding protein, did not affect spine-RNA granule distance. However, spines with a higher level of DBN1 exhibited a shorter distance to the nearest RNA granule down to a minimum of around 0.5 μm (Fig. 2d), and a DBN1 mutant that lacks the IDR strongly altered the observed vicinity between the spine and the RNA granule (Fig. 2d). In all, these observations point to a role of DBN1 in RNA granule recruitment at the base of spines.

**Fig. 2.**
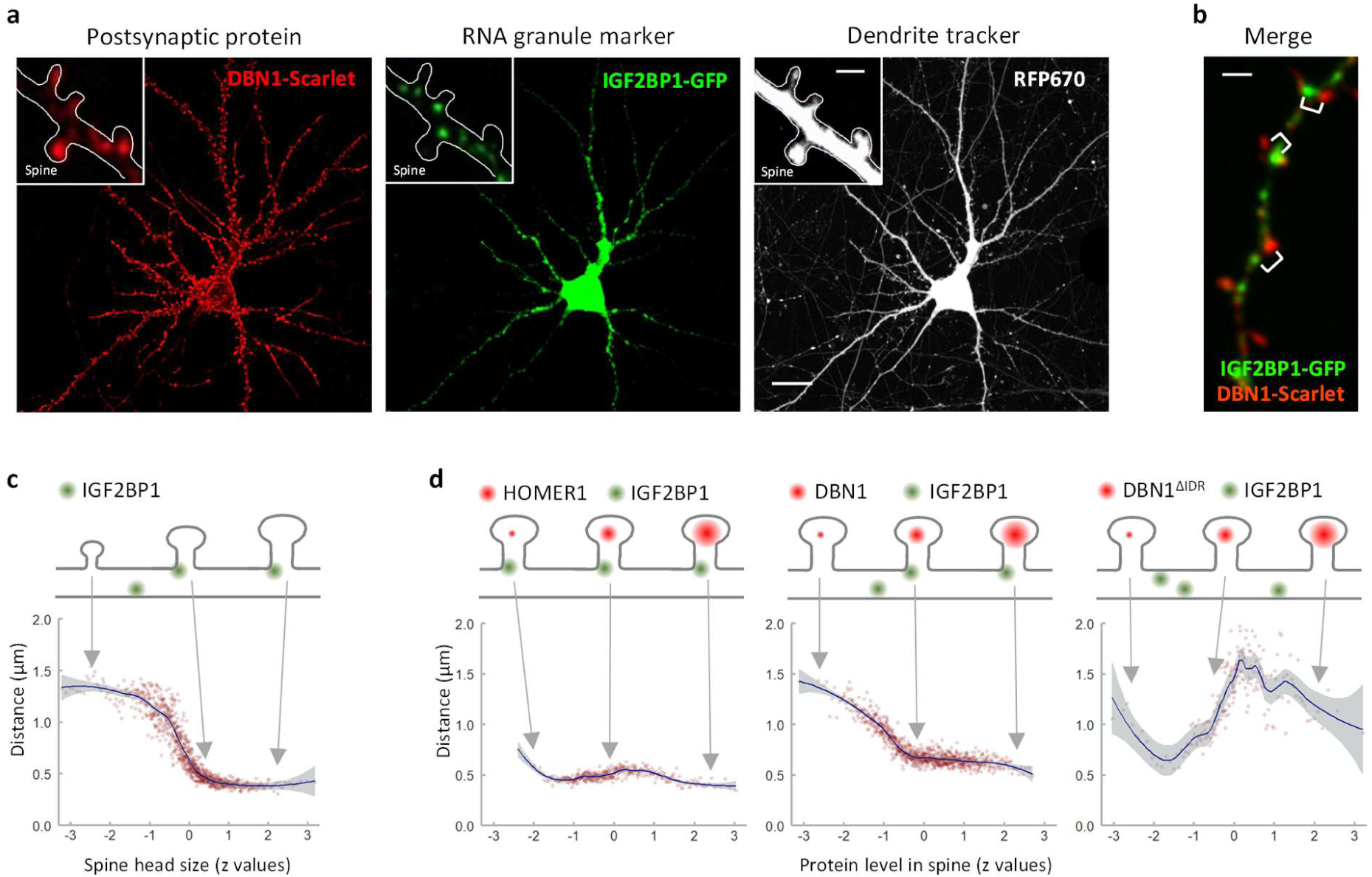
Correlated distribution of postsynaptic spines and RNA granules in dendrites. **a**, A representative hippocampal neuron expressing DBN1-Scarlet, IGF2BP1-GFP and RFP670 as a dendritic tracker (bar, 20 µm). Insets show a magnified dendritic segment (bar, 1 µm). **b**, A representative dendritic segment (bar, 1 µm) showing the distances determined in the distribution analysis. **c**, Spine head to RNA granule distance as a function of spine head size. **d**, Spine head to RNA granule distance as a function of the level of the indicated proteins in the spine head.

We also wanted to assess a possible correlation in size between the RNA granule and postsynaptic spine. To this end, we reanalysed our data and found that the size of RNA granules does not change in a range of small spines (Extended Data Fig. 2d). However, RNA granule size and spine head size displayed a strong positive correlation in the medium-to-large spine range, which points to the existence of positive feed-back mechanisms coordinating the postsynaptic compartment and RNA granule.

### Postsynaptic and RNA granule remodelling

The observed correlations in the dendritic distribution of postsynaptic spines-and RNA granules prompted us to analyse changes in protein proximity within either subcellular entity during synaptic activation. Thus, we expressed APEX2 fused to DBN1 or IGF2BP1 from lentiviral vectors in cortical neurons to perform biotinylation assays at 17DIV under basal, cLTP and post-cLTP conditions as mentioned above (Extended Data Fig. 1a). The DBN1-APEX2 and IGF2BP1-APEX2 fusions localised to dendrites as expected (Extended Data Fig. 3a) and attained levels comparable to the endogenous proteins (Extended Data Fig. 3b). A representative gel showing proteins biotinylated by the different APEX2 fusions under basal conditions is shown (Extended Data Fig. 3c). In these experiments, protein levels in streptavidin-pulldowns were made relative to those obtained from neurons expressing free cytoplasmic APEX2 as control. Thus, protein accessibility changes were corrected so that resulting scores would essentially measure the proximity of target proteins to either DBN1 or IGF2BP1 (Fig. 3a). As expected, actin-binding proteins and RNA granule protein components yielded high proximity scores to DBN1 and IGF2BP1, respectively, under basal conditions (Fig. 3b). In addition, described physical interactors were significantly enriched among proteins with high proximity to either DBN1 or IGF2BP1 (Extended Data Fig. 3d). In all, these data support the validity of our experimental design.

**Fig. 3.**
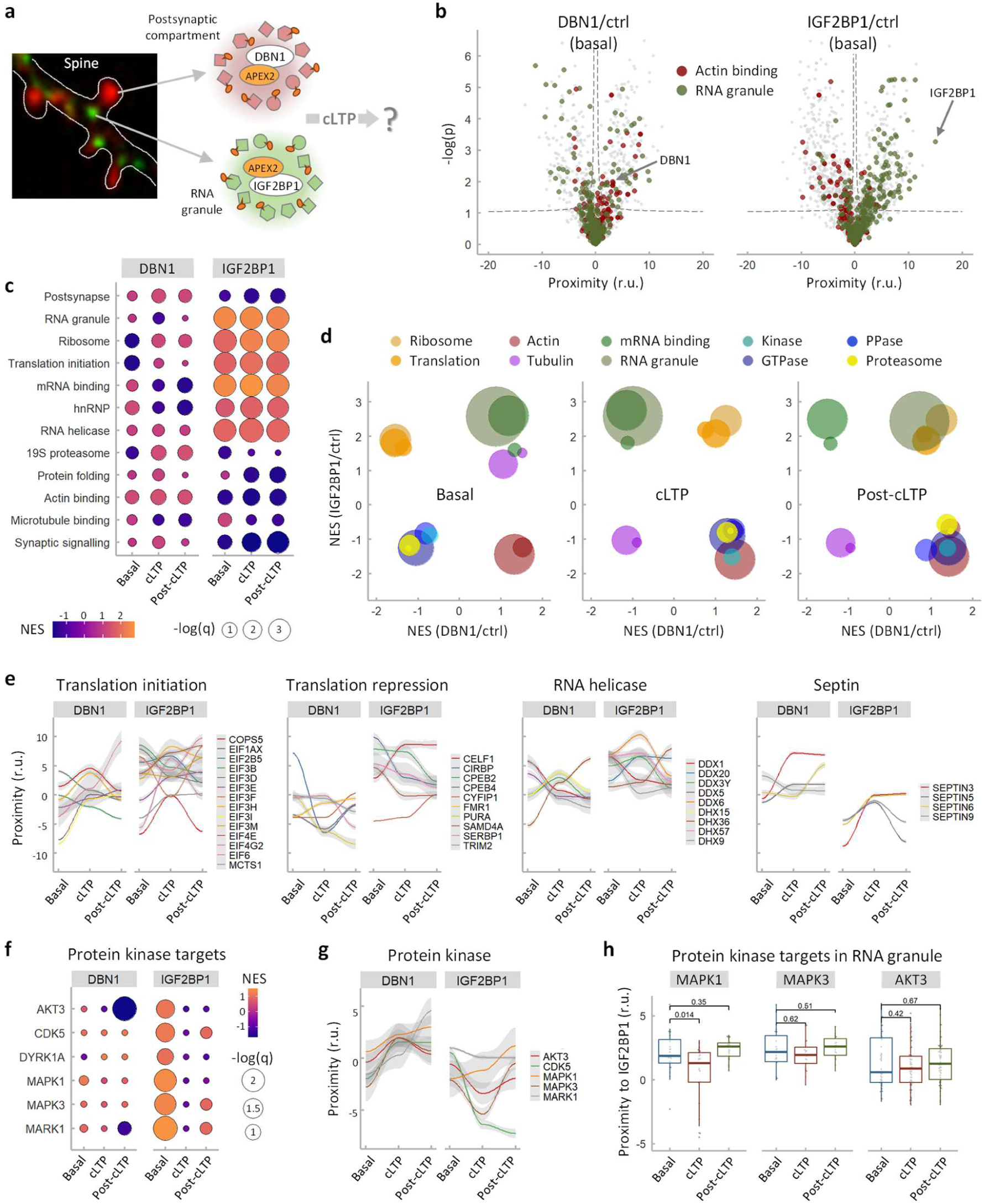
Protein-proximity changes in the postsynaptic compartment and RNA granule by cLTP. **a**, Schematic of the experimental approach to analyse changes in protein proximity in cortical neurons during cLTP. **b**, Volcano plots with proximity scores of individual proteins to DBN1-APEX2 or IGF2BP1-APEX2 (red and green dots represent proteins belonging to actin-binding or RNA-granule protein sets, respectively). **c**, GSEA results for the indicated selection of protein sets. **d**, Enrichment scores of the indicated protein sets by proximity to DBN1-APEX2 and IGF2BP1-APEX2 under basal, cLTP and post-cLTP conditions (bubble size indicates protein number in each protein set). **e**, DBN1-APEX2 and IGF2BP1-APEX2 proximity scores of the indicated proteins during cLTP. **f**, GSEA results for the indicated selection of protein kinase targets. **g**, DBN1-APEX2 and IGF2BP1-APEX2 proximity scores of the indicated protein kinases during cLTP. **h**, Proximity scores corresponding to the indicated protein kinase targets under basal, cLTP and post-cLTP conditions.

Under basal conditions, ribosomal proteins, translation initiation factors, protein kinases and GTPases were strongly underrepresented in the DBN1-proximal subproteome of cortical neurons. However, synaptic stimulation by cLTP completely inverted the normalised enrichment score (NES) of these protein sets (Fig. 3c,d) and caused a very significant increase in their proximity to DBN1 (Extended Data Fig. 3e). Notably, the increased proximity of these proteins to DBN1 was sustained during the post-cLTP period. Hacisuleyman *et al*.^21^ used a PSD95-TurboID fusion to biotinylate proximal ribosomes and analyse the postsynaptic translatome under basal and post-KCl conditions. Giving support to our observations, a reanalysis of these datasets resulted in high enrichment scores for ribosomal proteins and GTPases among proteins that increased their proximity to the PSD after KCl stimulation (Extended Data Fig. 3f).

Other protein categories inverted their NES in the DBN1 dataset under cLTP and post-cLTP conditions. The 19S proteasome displayed an increased and sustained enrichment in the vicinity of DBN1 after stimulation (Fig. 3c,d). Of note, the 19S regulatory subunit of the proteasome has been recently shown to be involved in regulating ubiquitin-dependent protein interactions in the postsynaptic compartment^31^. In contrast, RNA binding proteins -including hnRNPs-exhibited an opposite behaviour to that shown by the ribosome and translation factors, becoming more underrepresented in the proximity of DBN1 (Fig. 3c,d).

Protein categories in the IGF2BP1-proximal subproteome were more resistant to cLTP. The ribosome and translation initiation factors were already enriched among proteins with high proximity scores in the IGF2BP1 dataset under basal conditions (Fig. 3c,d) and did not increase their NES further by cLTP. Accordingly, their proximity scores did not change significantly either (Extended Data Fig. 3e).

Finally, regardless of their corresponding categorical enrichment, some individual proteins displayed strong fluctuations in their proximity to DBN1 and/or IGF2BP1 when comparing basal to cLTP and post-cLTP conditions (Fig. 3e). Among them, translation initiation factor EIF4G2 exhibited a large increase in proximity to both DBN1 and IGF2BP1 in the post-cLTP state. This translation initiation factor participates in a cap binding protein complex that acts in cap-dependent and independent translation. More specifically, EIF4G2 binding at the 5′ UTR of neuritic mRNAs has been associated with enhanced translation in response to synaptic activation by KCl and DHPG^21^. DHX36 proximity to both DBN1 and IGF2BP1 also increased under cLTP and post-cLTP conditions. This RNA helicase has been found in RNA granules^15^ and loss of DHX36 leads to an accumulation of translationally inactive target mRNAs in stress granules^32^. DDX3Y, another helicase with increased proximity to DBN1 and IGF2BP1 upon cLTP, has an X-linked paralog that attenuates RNA-RNA interactions and facilitates RNP granule dissolution^33^. DDX6 was only detected in the vicinity of IGF2BP1, but peaked to maximal levels upon cLTP. This RNA helicase has been shown to be involved in RNA granule assembly and disassembly during neuronal development and ageing^34–36^. SEPT3 also exhibited a higher proximity to both DBN1 and IGF2BP1 upon cLTP. Interestingly, septins form ring-shaped structures at the neck and base of dendritic spines and, in hippocampal neurons, play a key role in synaptic plasticity. In particular, SEPT3 interacts with myosin and promotes ER entry into spines in a calcium-dependent manner^37^.

EIF3D, CIRBP, and SERBP1 displayed an opposite behaviour and decreased their proximity to DBN1 and IGF2BP1 under cLTP and post-cLTP conditions. EIF3D, a factor that mediates translation initiation of a subset of mRNAs in proliferating cells^38^, has been shown to interact with the ribosome and G3BP2 ^15^, suggesting a direct role of EIF3D in ribosome recruitment into RNA granules. On the other hand, whereas CIRBP has been found in stress granules acting as a translational repressor^39^, SERBP1 is phosphorylated by TORC1 in dormant ribosomes to relieve translation inhibition ^40^.

Based on current phosphosite databases^41^, we selected target proteins of synaptic kinases and analysed their proximity to both DBN1 and IGF2BP1 during cLTP. To facilitate gene enrichment analysis, we first used the large dataset from *in vitro* kinase assays. No significant enrichment of kinase target proteins was observed when considering their proximity to DBN1 under basal, cLTP or post-cLTP conditions. In contrast, targets of many synaptic kinases were found to be strongly enriched in the vicinity of IGF2BP1 under basal conditions and, interestingly, six of them lost their enrichment upon cLTP (Fig. 3f). Regarding the six protein kinases themselves, all gained proximity to DBN1 during cLTP but only one of them, MAPK1, also increased its proximity to IGF2BP1 (Fig. 3g). Finally, we focused on target proteins of the RNA granule and used phosphosite datasets validated in vivo in perturbation experiments^41^. Again, only MAPK1-target proteins displayed a significant decrease in their proximity to IGF2BP1 upon cLTP (Fig. 3h), suggesting a role of this important signalling kinase in modulating the disassembly of some components of the RNA granule after synaptic activation.

### Postsynaptic cues sustain the RNA granule

Our proximity-labelling approach unveiled specific common features in the remodelling of the postsynaptic compartment and RNA granule during and after synaptic stimulation, suggesting the existence of signalling mechanisms linking both subcellular entities. DBN1 is essential for proper dendritic spine development and postsynaptic protein accumulation^42^. In addition, we found that expression of a DBN1 mutant lacking the IDR imposed a dominant perturbation of the distribution of RNA granules along dendrites (Fig. 2c). Thus, we hypothesised that this actin-binding protein could contribute to the signalling mechanisms linking the postsynaptic compartment and RNA granule. To test this idea, we downregulated DBN1 levels in cortical neurons expressing APEX2 or IGF2BP1-APEX2 from lentiviral vectors and performed biotinylation assays at 17DIV under basal, cLTP and post-cLTP conditions as above (Fig. 4a, Extended Data Fig. 1a). Under our experimental conditions, Dbn1 expression decreased more than 5-fold (Extended Data Fig. 4a) and the number of dendritic spines per μm suffered a strong reduction (Extended Data Fig. 4b,c) as described ^42^. Dbn1 downregulation did not affect the number of dendritic IGF2BP1-RNA granules per μm but produced a significant, albeit small, increase in the RNA granule-spine distance (Extended Data Fig. 4d,e).

**Fig. 4.**
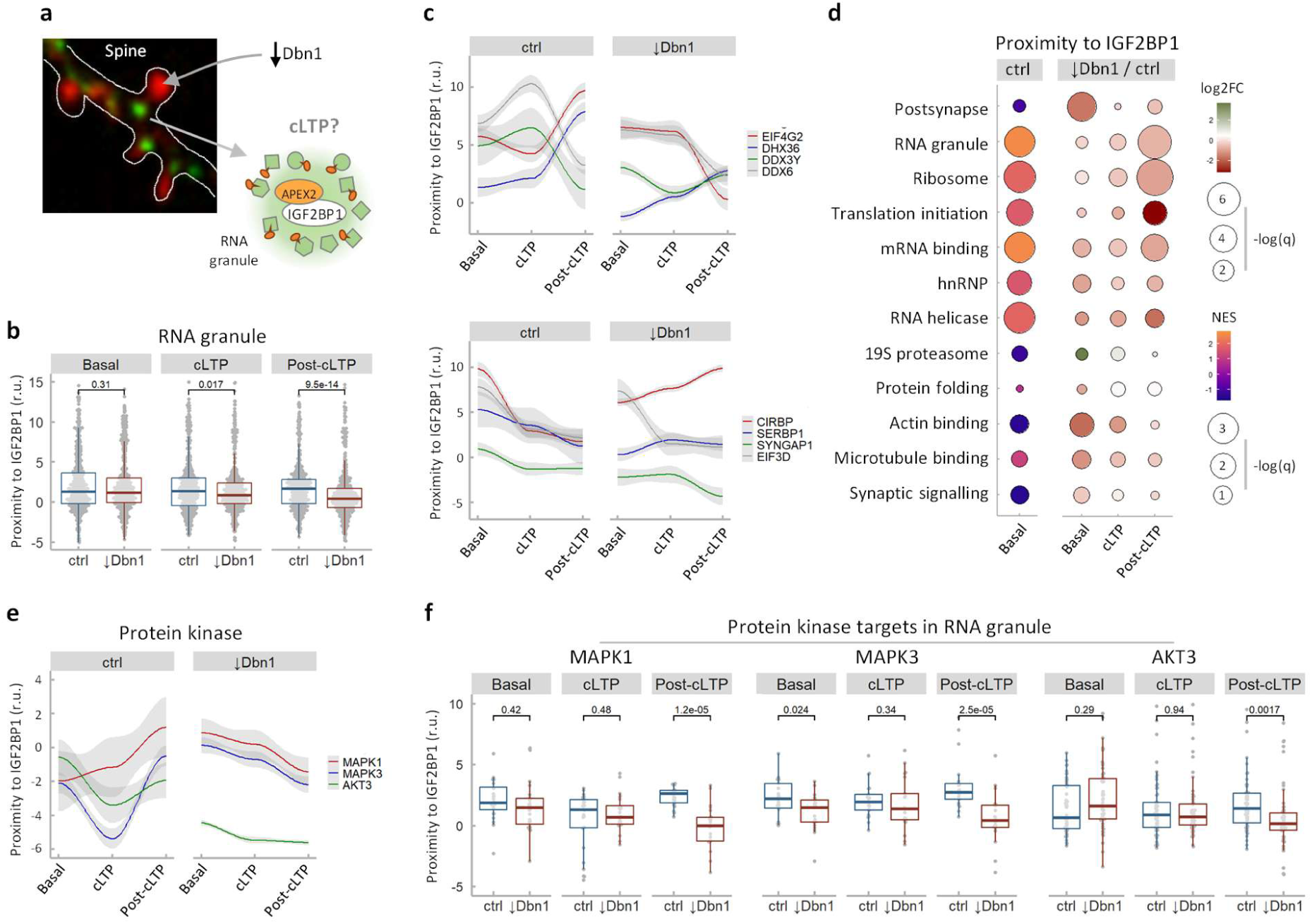
Postsynaptic cues sustain the RNA granule upon cLTP. **a**, Schematic of the experimental approach to analyse changes in protein proximity to IGF2BP1-APEX2 after Dbn1 downregulation in cortical neurons during cLTP. **b**, IGF2BP1-APEX2 proximity scores corresponding to the RNA granule protein set under basal, cLTP and post-cLTP conditions in control (ctrl) or Dbn1-knockdown (↓Dbn1) corical neurons. **c**, IGF2BP1-APEX2 proximity scores of the indicated individual proteins as in (b). **d**, GSEA results and proximity changes (↓Dbn1/ctrl) in Dbn1-knockdown vs control cortical neurons for the indicated selection of protein sets. **e**, IGF2BP1-APEX2 proximity scores of the indicated individual proteins as in (b). **f**, IGF2BP1-APEX2 proximity scores corresponding to the indicated protein kinase target sets under basal, cLTP and post-cLTP conditions.

Dbn1 downregulation did not affect the proximity of RNA granule proteins to IGF2BP1 in basal samples but did cause an overall 3-fold decrease in their proximity score under post-cLTP conditions (Fig. 4b). Proximity of ribosomal proteins, translation initiation factors and mRNA binding proteins to the RNA granule reporter exhibited a very similar dependence on DBN1 (Extended Data Fig. 4f), again particularly pronounced in the post-cLTP samples. In contrast, proximity of glycolysis enzymes was unaffected by Dbn1 downregulation. With the exception of EIF3D, Dbn1 downregulation imposed strong changes in the proximity to IGF2BP1 during cLTP when considering translation factors and RNA helicases already highlighted in the previous section by their special functional relevance, namely EIF4G2, DHX36, DDX3Y, CIRBP and SERBP3 (Fig. 4c). SYNGAP1, an abundant postsynaptic GTPase activator, displayed a similar proximity value upon cLTP, but dropped more than 5-fold under both basal and post-cLTP conditions. When considering the whole IGF2BP1-proximal subproteome, Dbn1 knockdown caused a stronger loss of proximity to IGF2BP1 in those protein categories displaying a higher proximity in the basal state, i.e. RNA granule, ribosome, translation initiation, mRNA binding and RNA helicase (Fig. 4d).

Finally, Dbn1 downregulation also affected the proximity of MAPK1, MAPK3 and AKT3 to IGF2BP1, and these three synaptic protein kinases decreased their proximity score mostly in post-cLTP samples (Fig. 4e). Notably, the respective target proteins of the RNA granule also displayed a strong decrease in proximity to IGF2BP1 under post-cLTP conditions (Fig. 4f). In all, our data support the notion that postsynaptic signals play a relevant role in the mechanisms that maintain the composition of RNA granules after synaptic stimulation by cLTP.

## Discussion

Synaptic plasticity relies on local translation of dendritic transcripts that are transported as RNA granules from the soma to synapses. Although it is generally assumed that signals emanating from the postsynaptic compartment play a major role in nearby recruitment and partial disassembly of RNA granules^28,43,44^, the underlying mechanisms are still a mystery. Here we present protein accessibility and proximity assays focusing on the postsynaptic compartment and RNA granule subproteomes to analyse the effects of synaptic activation by cLTP on their composition and interdependence. First, we observed extensive alterations in the accessibility of RNA granule proteins in the neuritic cytoplasm upon and after cLTP, particularly affecting the ribosome and translation initiation factors. After a detailed analysis of the increase in biotinylation of ribosomal proteins as a function of the number of tyrosines exposed or hidden in the 80S structure, we concluded that the gain in accessibility by cLTP is not due to dissociation of the ribosome into single components but, rather, to the withdrawal of ribosomes from large assemblies such as the RNA granule. In support of this idea, hnRNPs and other mRNA binding proteins found in the RNA granule also displayed a sharp increase in their accessibility upon cLTP. Interestingly, most of these changes in accessibility were also observed in the post-cLTP sample, suggesting a role in synaptic plasticity. Regarding the postsynaptic compartment, we observed many signalling proteins displaying striking changes in their accessibility upon cLTP, supporting the idea that synaptic activity drives a profound remodelling of signalling pathways in the PSD^45^.

To study the effects of cLTP on the composition of the RNA granule and postsynaptic compartment, we used APEX2 fusions to IGF2BP1 and DBN1 as respective baits to analyse changes in protein proximity during synaptic activation. Whereas the proximal subproteome of IGF2BP1 was more resistant, the postsynaptic compartment displayed profound alterations during cLTP, which particularly affected the translational machinery. Indeed, the ribosome and translation initiation factors strongly increased their presence in the vicinity of DBN1, thus favouring their engagement in local translation.

Some ribosomal proteins are differentially up- or downregulated in the nascent proteome of axons in response to distinct cues^46^. We found that not all ribosomal proteins displayed the same increase in proximity to DBN1 in response to cLTP, raising the possibility that postsynaptic signals remodel the ribosome in different ways to regulate translation of specific mRNAs^47,48^.

Contrary to the ribosome and translation initiation factors, RNA binding proteins decreased their proximity to the postsynaptic compartment, indicating that not all components of the RNA granule are relocated similarly by cLTP. Considering the role of hnRNPs in RNA granule condensation^44^, their increased underrepresentation in the vicinity of DBN1 upon cLTP would suggest that mRNAs increase their availability as translationally-competent in the postsynaptic spine. In *X. laevis*, hnRNPs bind to both ribosomes and mRNAs in a translationally repressed state under basal conditions but, upon stimulation, dissociate from mRNAs and allow their translation in the growth cone^49^.

Supporting the view that synaptic activation uses different mechanisms to favour mRNA translation, we have identified several proteins that sharply increased their proximity to DBN1 or IGF2BP1 during cLTP. EIF4G2 has been shown to be involved in activating translation in response to synaptic stimulation by KCl and DHPG^21^. DHX36 is an RNA helicase found in RNA granules^15^ that is essential to relieve mRNAs from translational inhibition ^32^. DDX3Y has an X-linked helicase paralog that triggers RNP granule dissolution ^33^. Finally, DDX6 has been shown to be involved in RNA granule assembly and disassembly during neuronal development and ageing^34–36^. In contrast, other proteins (EIF3D, CIRBP and SERBP1) associated with translational repression ^15,38–40^, displayed an opposite behaviour and decreased their proximity to DBN1 and IGF2BP1 upon cLTP.

It is widely assumed that postsynaptic mechanisms remodel the RNA granule to stimulate local translation ^28,44^. Initial evidence based on fluorescence and electron microscopy revealed that neuronal RNA granules become less compact after KCl depolarisation ^19^. Later, using single mRNA-reporter systems, cLTP was shown to cause transient unmasking, motility, and local translation of paradigmatic mRNAs ^17,43,50^. Early work associated β-actin mRNA translation with SRC-dependent phosphorylation of IGF2BP1 ^51^ and, more recently, phosphorylation in two disordered linkers of IGF2BP1 has been shown to modulate RNP condensation ^52^. However, whether phosphorylation is a general mechanism to modulate RNA granule composition in the postsynaptic neuron is still a matter of study. Our proximity labelling experiments uncovered a key difference in the composition of the RNA granule and the postsynaptic compartment. Although the PSD is assumed to work as a signalling hub, proteins targeted by synaptic kinases are not particularly enriched in the postsynaptic compartment. Instead, they dominate the vicinity of the RNA granule reporter and, for some paradigmatic kinases, their enrichment was totally abolished by cLTP. These protein kinases displayed an increased proximity to DBN1 during cLTP, but only one of them, MAPK1, also showed increased proximity to the RNA granule after synaptic stimulation. Notably, MAPK1 targets decreased their proximity to the RNA granule reporter transiently upon cLTP. In all, these observations strongly support the notion that MAPK1 acts as a key cLTP-signalling kinase to phosphorylate RNA granule components and drive disassembly of ribosomes and translation initiation factors, thus triggering local translation of released mRNAs in the juxtaposed postsynaptic compartment. Finally, DBN1 downregulation caused altered proximity patterns in the vicinity of IGF2BP1 that also affected MAPK1, MAPK3 and AKT3 kinases and the corresponding targets in the RNA granule. These alterations were particularly important in post-cLTP samples, indicating that proper signalling mechanisms in the postsynaptic compartment play a key role in sustaining the molecular composition of the RNA granule.

## Acknowledgments

We thank L. Luna for technical assistance and C. Rose for editing the manuscript. We also thank K. Roy for shRNA constructs and B. Poquet for stimulating comments and suggestions. This work was funded by a grant from the Ministry of Science and Innovation of Spain and the European Union (FEDER) (PID2023-146803NB-I00) to C.G.

## Author contributions

C.G. and M.A. conceived the project. M.B.M., K.A., G.Y. and A.S. performed experiments. M.B.M., C.G. and M.A. performed data analysis. C.G. and M.A. guided experiments and data analysis. C.G. and M.A. wrote the manuscript with input from all of the authors.

## Methods

### Cell culture

We used hippocampal neurons for imaging experiments in which mature neurons with well-defined dendritic spines for morphological analysis were required, whereas cortical neuron cultures were used for proximity labelling experiments that required a large amount of initial cellular material.

Hippocampi and cortex were dissected from E17.5 embryos of Swiss mice (undetermined sex) in HBSS containing 0.6% glucose and 10 mM HEPES. After dissection, tissues were digested with 0.05% trypsin (Fisher, 15090046) and 0.02% EDTA (Merck-Sigma, E9884) in HBSS at 37°C for 15 min. Enzymatic digestion was stopped by washing the tissue three times with MEM (Fisher, 31095029) supplemented with 10% FBS (Biowest, S1810-500) and 0.6% glucose. Hippocampi were left to sediment between washes, and centrifugation was avoided to keep cell viability. Trypsin-treated tissue was then mechanically disaggregated by passing through a flame-polished Pasteur pipette (∼10 times). Cells were plated at desired density on Poly-D-lysine (Merck-Sigma, P7886)-coated plates (0.5 mg/ml Poly-D-lysine in borate buffer, pH 8.5) and maintained in MEM, 10% FBS and 0.6% glucose. After 2–4 h plating medium was substituted by neuronal maintaining medium: Neurobasal (Fisher, 11570556) medum supplemented with 2% B27 (Fisher, 11530536) and 1% GlutaMAX (Fisher, 35050-038). Primary hippocampal cultures were transfected using CalPhos mammalian transfection kit (Clontech, 631312) as previously described^1^. HEK293T cells (ATCC) were cultured in DMEM medium supplemented with 10% FBS. HEK293T transfection was carried out with Lipofectamine 2000 (Invitrogen, 11668030), according to the manufacturer’s instructions. Cells were routinely tested for mycoplasma contamination. Animal experimental procedures were approved by the ethics committee of the National Research Council of Spain (Approval number: B9900083).

### DNA vectors

Lentiviral constructs pLKO-RFP-shCtrl (Addgene, 69040) and pLKO-RFP-shDbn1 containing the shRNA sequence GCAGTCTATCTTTGGTGACCA (TRCN0000090077) against the Dbn1 gene were used in knockdown experiments. pFUW-FLAG-APEX2, pFUW-FLAG-APEX2-DBN1 and pFUW-FLAG-APEX2-IGF2BP1 were used to perform accessibility and proximity assays, and derived from pcDNA3 APEX2-NES (Addgene, 49386) and pFUW (Addgene, 14882). Dbn1, Dbn1^ΔIDR^ and Igf2bp1 were amplified using Phusion High-Fidelity DNA Polymerase (Thermo Fisher, 10024537) from mouse cDNAs. mEGFP-C3 (Clontech) and mScarlet-C1 (Addgene, 85044) were used as host vectors. GFP-Homer1b was a gift from M. Bosch. piRFP670-N1 (Addgene 45457) was used as a tracker. Primers used in all these cloning experiments are shown in Supplementary Table 1.

### Lentivirus production and infection

HEK293T cells were transfected with pFUW lentiviral vectors, envelope plasmid pVSV-G (Addgene, 8454) and packaging plasmid pCMV-dR8.2 dvpr (Addgene 8455). After 4 hours of transfection with Lipofectamine 2000, the transfection medium was replaced with 10 ml of neuronal maintaining medium. Lentiviruses were harvested 48h and 72h after transfection and passed through a 0.45 μm syringe filter (Fisher, 15216869). Supernatants were used to transduce primary cortical neurons between 4 and 7 DIV by replacing half of their media with medium containing lentiviral particles. Medium was completely changed by fresh neuronal maintaining medium three hours post-infection, and cortical neurons were cultured until 17 DIV.

### APEX2 biotinylation assays

Primary cortical neurons were plated in two 100 mm plates per condition at 4×10^6^ cells/plate, and infected at 7 DIV using pFUW-FLAG-APEX2, pFUW-FLAG-APEX2-DBN1 or pFUW-FLAG-APEX2-IGF2BP1 plasmids. In Dbn1 knockdown experiments, neurons were co-infected with pLKO-RFP-shDbn1 and pFUW-FLAG-APEX2 or pFUW-FLAG-APEX2-IGF2BP1. At 17 DIV, prior to APEX2 activation, neurons were incubated with 500 μM biotin-phenol (BP) (Merck-Sigma, SML2135) for 30 min at 37°C. APEX2 labelling was performed by adding hydrogen peroxide (Merck-Sigma, 88597) to a final concentration of 1 mM for 1 min and the biotinylation reaction was immediately stopped by washing three times with STOP/Quench buffer: 5 mM Trolox ((+/-)-6-Hydroxy-2,5,7,8-tetramethylchromane-2-carboxylic acid) (Merck-Sigma, 238813), 10 mM sodium ascorbate (Merck-Sigma, A7631), 10 mM sodium azide (Merck-Sigma, S2002), 1 mM CaCl_2_ and 0.5 mM MgCl_2_ in phosphate-buffered saline (PBS) at pH 7.4. Samples were collected using cell scrappers and suspended in cold lysis buffer (4M urea, 50 mM Tris pH 8, 150 mM NaCl, 5 mM EDTA, 0.5 % sodium deoxycholate, 0.1 % SDS, 1 % Triton X-100, 5 mM Trolox, 10 mM sodium ascorbate and 10 mM sodium azide). To ensure complete cell lysis, samples were passed repeatedly through a syringe needle (26G). Then, one volume of 55 % trichloroacetic acid (Merck-Sigma, T6399) was added and samples were incubated on ice for 15 min. Proteins were precipitated at 14000 rpm at 4 °C for 10 min. Pellets were washed with -20°C cold acetone (Merck-Sigma, 179124) three times. Dry pellets were resuspended in cell lysis solution (8M urea, 100 mM sodium phosphate pH 8, 1% SDS, 100 mM NaHCO3 and 10 mM TCEP) until they were completely dissolved. The resuspended protein solutions were centrifuged at 14000 rpm at room temperature for 10 min and supernatants were transferred to new microcentrifuge tubes. Biotinylated proteins were captured onto streptavidin magnetic beads (Fisher, 10150874). Magnetic beads (75 µl per condition) were washed three times in 4M urea and 0.5 % SDS and added to each sample. Beads were incubated overnight at 4 °C and then separated from solutions by using a magnetic rack, washed once with 4M urea and 0.5 % SDS and three times with 50 mM Tris pH 8. The beads were frozen at -80°C prior to on-bead digestion for MS analysis. Biotinylation was monitored by immunoblotting using a streptavidin-HRP conjugate antibody as described below.

For cLTP stimulation, cortical neurons were pre-incubated in Mg^2+^-free artificial cerebrospinal fluid (ACSF) with 0.5 μM TTX (Tocris, 1069), 1 μM strychnine (Merck-Sigma, S0532) and 20 μM bicuculline methiodide (Merck-Sigma, 14343) for 30 min followed by stimulation of synaptic N-methyl-D-aspartate receptors by addition of 200 µM glycine (Tocris, 0219) as co-agonist for 10 min. For post-cLTP samples, the medium was aspirated and cortical neurons were incubated in Mg^2+^-free ACSF for 30 min.

### Immunofluorescence

Primary hippocampal neurons were plated into 4-well plates containing coverslips (Fisher, 10507591) at a density of 2×10^4^ cells/cm^2^, infected at 7 DIV with the indicated lentivirus, and fixed at 14 DIV with 4 % paraformaldehyde (Merck-Sigma, P6148) with 4 % sucrose at 4°C for 20 min. After fixation, cells were washed with PBS and permeabilized using 0.1 % Triton-X (Merck-Sigma, 11332481001) in PBS at 4°C for 5 min. Non-specific binding sites were blocked with 5 % normal goat serum (Fisher, PCN5000) in PBS for 30 min at room temperature. After blocking, neurons were incubated overnight at 4°C with rabbit anti-FLAG (1:1000, Merck-Sigma, F7425) and mouse monoclonal anti-MAP2 (1:1000, Merck-Sigma, M1406) primary antibodies. Donkey anti-rabbit Alexa Fluor™ 568 (1:1000, Fisher-Invitrogen, A10042) and goat anti-mouse Alexa Fluor™ 488 (1:1000, Fisher-Invitrogen, A-11001) secondary antibodies were incubated for 2h at room temperature. Image stacks were acquired in an Andor Dragonfly 505 confocal microscope under 63x 1.4-NA oil-immersion objective and processed using ImageJ for visualization.

### Live-imaging

For live imaging experiments, primary hippocampal neurons were plated at a density of 2×10^4^ cells/cm^2^ in Ibidi μ-Dish 35 mm (Inycom, 81156). Neurons were co-transfected at 7 DIV with piRFP670-N1 and plasmids expressing fusions of DBN1, IGF2BP1 and HOMER1 to mScarlet or mGFP, as indicated. In Dbn1 knockdown experiments, cells were co-transfected with plasmids expressing pLKO-RFP-shCtrl or pLKO-RFP-shDbn1 and GFP-IGF2BP1. At 14 DIV, neurons were live-imaged to determine the localization and levels of DBN1 and IGF2BP1 through dendrites. Images were taken using an Andor Dragonfly 505 spinning-disk microscope equipped with a 5 % CO2, 37°C humidified chamber. Stacks (1 μm z-interval) were taken under a 63x 1.4-NA oil-immersion objective. Laser power and all other acquisition parameters were kept constant throughout images and conditions. To assist neurite tracking, neurons co-expressed RFP670, a far-red fluorescent protein. Individual spines were identified in neurons expressing mScarlet-DBN1 or GFP-HOMER1, and their fluorescence levels measured. Tracker fluorescence in these protrusions was also measured to normalize the synaptic fluorescence signal and obtain protein concentration in relative units. In addition, RNA granules were detected in neurons expressing mScarlet-or mGFP-IGF2BP1. Foci positions were determined and distances to the closest RNA granule were determined. Single protrusion and focus data were binned (n=100) and plotted with the corresponding trend line and confidence limits.

### Immunoblotting

Primary cortical neurons were lysed in 1x SR (2 % SDS and 0.125M Tris-HCl pH 6.8) and sonicated. Following electrophoresis, proteins were transferred from the SDS-PAGE gel onto a nitrocellulose membrane (Merck, GE10600002) using a semi-dry transfer system (Bio-Rad, 1703940).

The membrane was blocked with non-fat dry milk at room temperature for 1h and incubated with mouse monoclonal anti-DBN1 (Fisher-Invitrogen, PIMA528543, 1:200), or rabbit anti-IGF2BP1 (IMP1) (MBL, RN007P, 1:200) primary antibodies for 1h. Anti-mouse IgG-HRP (Merck, GENXA931, 1:10000) and anti-rabbit IgG-HRP (Merck, GENA934, 1:10000) secondary antibodies or streptavidin-HRP (Fisher, N100, 1:1000) were incubated for 1h. Membranes were treated for 5 min with Clarity Western ECL substrate (Bio-Rad, 1705061), and images were captured in an Odissey M (LiquorBio) and analysed using ImageJ software.

### RNA extraction and real-time quantitative PCR

Primary cortical neurons were infected at 7 DIV with lentivirus pLKO-RFP-shCtrl and pLKO-RFP-shDbn1. At 14 DIV RNA was obtained using EZNA total RNA purification kit (Omega Bio-tek, R3634-01) following the manufacturer’s instructions. Then, samples were digested with ribonuclease-free deoxyribonuclease I (Roche, 11119915001) and 1 μg of RNA was reverse-transcribed into cDNA using the Maxima H Minus First Strand cDNA Synthesis Kit (Fisher, K1651). qPCR was performed with Taqman probes (6xFAM-BQ1) on a LightCycler@96 Real-Time PCR system (Roche) according to manufacturer’s instructions. qPCR oligonucleotides and probes are shown in Supplementary Table 2.

### APEX2 proteomic analysis

Proteins were subject to on-bead trypsinization and the resulting peptides were separated by reversed-phase chromatography for data-independent acquisition quantitative MS analysis in an Orbitrap Fusion LUMOS (BGI Genomics Co.). Proteins identified in all samples are detailed in Supplementary Table 3. Matched proteins were filtered for possible contaminants, and relative iBAQ data were log2 transformed. Overall, 3,645 proteins were detected. Missing signals were imputed by random sampling from normal distributions and, for missing replicates the minimal value in the dataset was increased 8-fold and used as mean. For every protein detected, an accessibility score was calculated as the log2 ratio of the relative iBAQ in streptavidin pulldown and input samples from cortical neurons expressing APEX2. In contrast, proximity scores were calculated as the log2 ratio of the relative iBAQ in streptavidin pulldowns from DBN1-APEX2 or IGF2BP1-APEX2 to the relative iBAQ from APEX2 expressing neurons.

### GSEA

Normalized enrichment scores were calculated using the GSEA software^2^. Proteins were ranked by their accessibility or proximity scores as indicated and preliminary analyses were done using the Molecular Signatures Database^3^. Specific gene sets shown in Fig. 1, Fig. 3 and Fig. 4 can be found in Supplementary Table 4.

### Statistical analysis

Pair comparisons were performed with the nonparametric Mann-Whitney test, and the resulting p values are shown in the corresponding figure panels. Data are displayed as median and quartile (Q) values. Tendency lines were obtained by non-parametric local regression with a confidence level of 0.9. Protein levels by immunoblotting and mRNA levels by RT-PCR were determined in triplicate samples, and mean ± SEM values were plotted.

### Software

SpineJ and DendFociJ are ImageJ (Wayne Rasband, NIH) plugins available with a basic set of instructions upon request.

### Data availability

The mass spectrometry proteomics data have been deposited to the ProteomeXchange Consortium through the PRIDE partner repository (PXD065056). All plasmidic and lentiviral constructs are available upon request.

---

**Extended Data Fig. 1.**
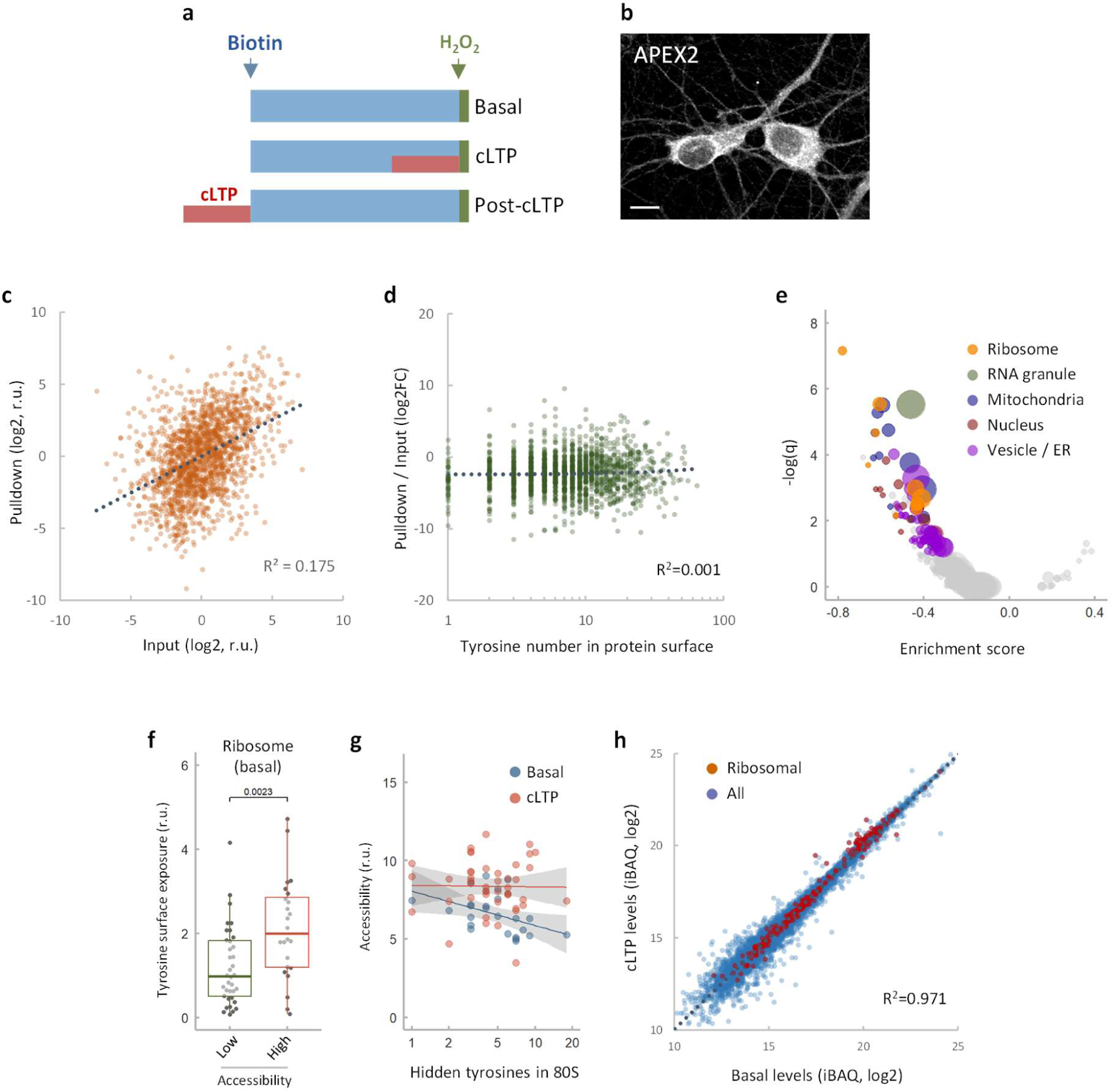
cLTP alters accessibility of postsynaptic and RNA granule proteins. **a**, Experimental design of biotinylation assays in cortical neurons under basal, cLTP and post-cLTP conditions. **b**, A representative αFLAG immunofluorescence image of cortical neurons expressing FLAG-APEX2 fused to a nuclear export signal sequence from a lentiviral vector. **c**, Proteomic analysis of streptavidin pulldown and input samples from cortical neurons subject to biotin labelling under basal conditions. **d**, Pulldown to input ratios as a function of the predicted number of surface tyrosines per protein. **e**, Volcano plot with enrichment scores for the indicated selection of protein sets (bubble size indicates protein number in each protein set). **f**, Tyrosine surface exposure in ribosomal proteins with low and high accessibility scores as determined by their streptavidin pulldown / input ratios. **g**, Accessibility scores corresponding to ribosomal proteins as a function of the number of surface tyrosines hidden in the 80S ribosome under basal (blue) and cLTP (orange) conditions. **h**, Proteomic analysis of cortical neurons under basal and cLTP conditions. Ribosomal proteins are indicated (red dots).

**Extended Data Fig. 2.**
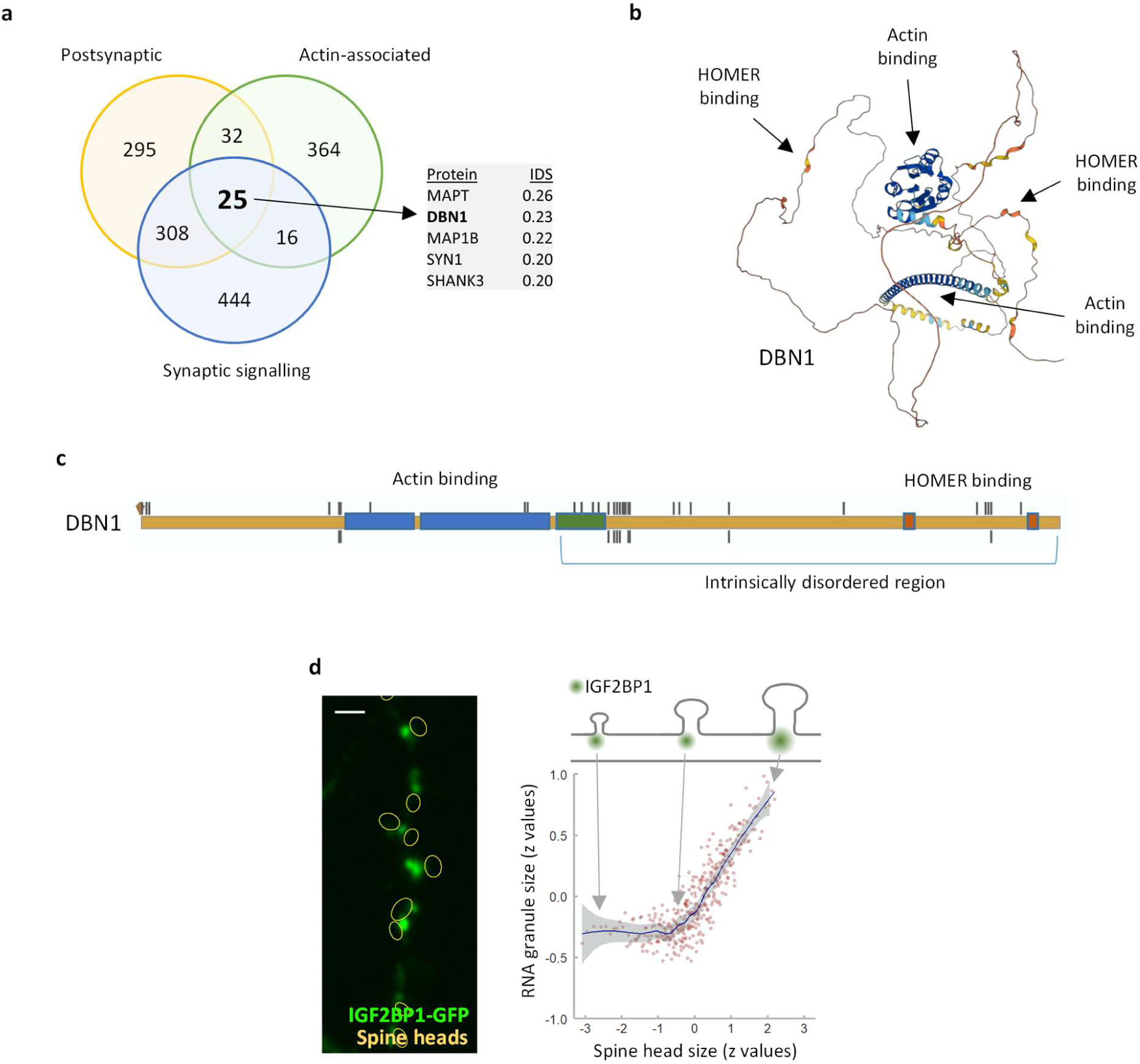
Correlated distribution of postsynaptic spines and RNA granules in dendrites. **a**, Venn diagram of postsynaptic actin-associated proteins that have been shown to be involved in signalling pathways, indicating those displaying the highest intrinsic disorder score (IDS). **b**, Alpha2Fold structural prediction of DBN1 showing the actin and HOMER binding domains. **c**, Schematic of DBN1 with phosphorylation residues in PhosphoSite. **d**, RNA granule size as a function of spine head size. A representative dendritic segment (bar, 1 µm) is shown at the left.

**Extended Data Fig. 3.**
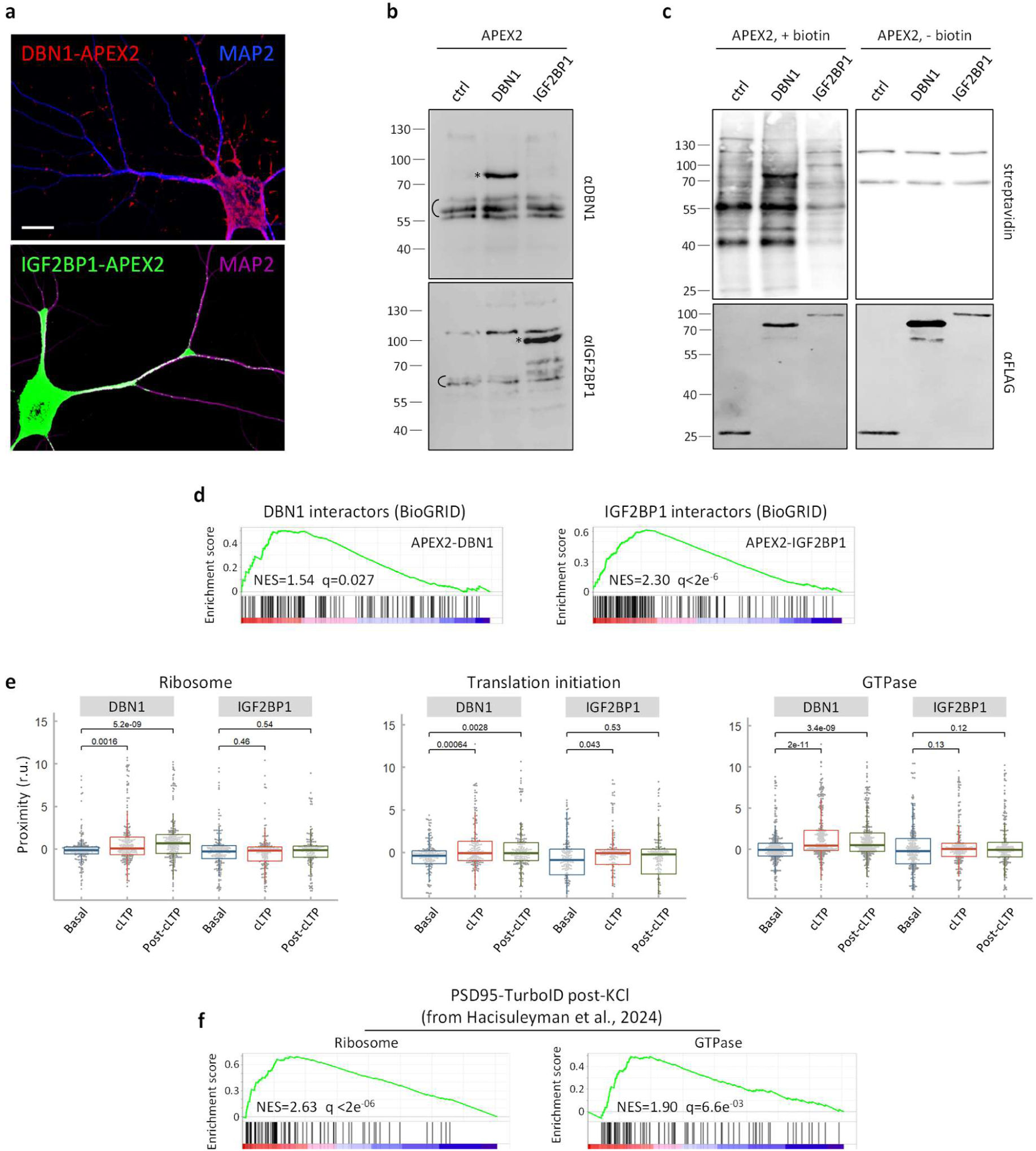
Protein-proximity changes in the postsynaptic compartment and the RNA granule by cLTP. **a**, Representative immunofluorescence images of cortical neurons expressing DBN1-APEX2 or IGF2BP1-APEX2. Map2 is shown as dendritic tracker (bar, 20 µm). **b**, Immunoblot analysis of cortical neurons infected with APEX2 (ctrl), APEX2-DBN1 or APEX2-IGF2BP1 lentivirus, showing endogenous (C) and APEX2-tagged (*) proteins. **c**, Proximity labelling assays of cortical neurons expressing APEX2 (ctrl), APEX2-DBN1 or APEX2-IGF2BP1 in the presence or absence of biotin as control. **d**, GSEA results of the APEX2-DBN1 or APEX2-IGF2BP1 proximity datasets. Barcode plots indicate the rank position of DBN1 and IGF2BP1 physical interactors (BioGRID). Normalized enrichment scores (NES) and the corresponding FDR values are shown. **e**, DBN1-APEX2 and IGF2BP1-APEX2 proximity scores corresponding to the indicated protein sets under basal, cLTP and post-cLTP conditions. **f**, GSEA results of the PSD95-TurboID proximity dataset from Hacisuleyman et al.^20^ Barcode plots indicate the rank position of ribosomal and GTPase proteins. Normalized enrichment scores (NES) and the corresponding FDR values are shown.

**Extended Data Fig. 4.**
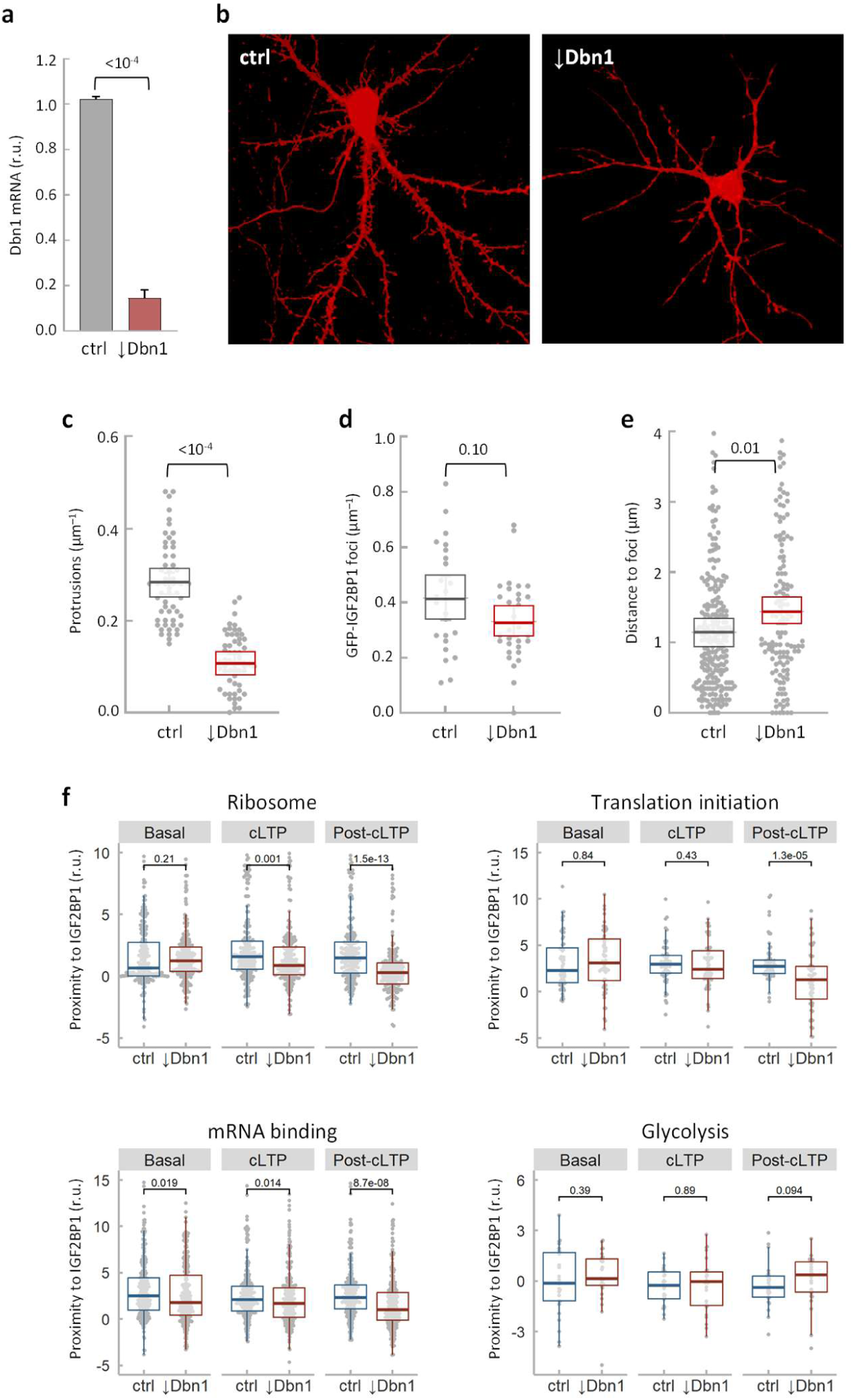
Postsynaptic cues sustain the RNA granule upon cLTP. **a**, Dbn1 mRNA levels in cortical neurons infected with control (ctrl) or shDbn1 (↓Dbn1) lenivirus expressing RFP as reporter. **b-e**, Representative images (b), protrusions per µm (c), IGF2BP1-Scarlet foci per µm (d) and spine head to IGF2BP1-Scarlet foci distances (e) from control (ctrl) and shDbn1 (↓Dbn1) hippocampal neurons. **f**, IGF2BP1-APEX2 proximity scores corresponding to the indicated protein sets from control (ctrl) and shDBN1 (↓Dbn1) corical neurons under basal, cLTP and post-cLTP conditions.

